# Quantitative analysis of the orientation of white stork nestlings’ parents from video monitoring

**DOI:** 10.1101/2021.09.11.459736

**Authors:** Ariane Gayout

**Affiliations:** Univ Lyon, ENS de Lyon, Univ Claude Bernard Lyon 1, CNRS, Laboratoire de Physique, F-69342 Lyon, France

**Keywords:** Behavioral correlations, *Ciconia ciconia*, citizen science, image processing, parental care, weather influence, webcam

## Abstract

Video-monitoring has become in the last decades common practice for animal observation and conservation purposes. In Ornithology, it is mostly used for tracking predators and nest surveillance, but, with the rapid development and spreading of webcams on nests for educational purposes, new opportunities arise for behavioral investigation, through citizen science for instance. In this article, we use video-monitoring from a public webcam on a White stork (*Ciconia ciconia*) nest and perform systematic image analysis to record the positioning and orientation of the guarding parent on the nest, during the nestling period over 60 days. From this data of 450 orientation samples, correlations with weather parameters are drawn. Our results suggest that the sun is responsible for most of the orientation with an hourly dependence, while the wind has prevalence during rainy days. A change in the parent behavior is also observed around the time the nestlings are known to attain their maximal weight. These preliminary findings provide new insights on weather influence on parental care behavior likely linked with the parent’s sensing. The versatility of the proposed method allows for behavioral studies on a wide variety of species.

## Introduction

In the midst of the recent developments of technology, Ornithology has obtained in the last decades strong tools to deepen our understanding of bird behavior. In particular, Global Positioning System (GPS) provides insights on complex migratory behaviors (Mandel et al. 2008, Kumar et al. 2020), while accelerometers help uncovering subtleties in raptor flight (Halsey et al. 2009, Laurent et al. 2021). Video monitoring has also become a tool for nest observation and conservation purposes, with predator identification for instance (Stake et al. 2004, Sabine et al. 2005, Bolton et al. 2007, Coates et al. 2008). Among the various technologies available for video monitoring, webcams, whose network has expanded drastically in the last decade, are particularly interesting for quantitative behavioral observations, thanks to their universality, the continuous flow of information and the possibility of citizen science (Schulwitz et al. 2018, Wood et al. 2020).

In this article, we use video monitoring on a White stork (*Ciconia ciconia*) nest to investigate parental behavior using image processing as a novel method to expand the scientific use of webcams. Technologies like GPS and nest platforms have already benefitted White storks (Tryjanowki et al. 2009). Well appreciated in the collective imagination and already subject to citizen science (Thabethe and Downs 2018), storks are also known to be great indicators for climate variations from their migratory behavior (Berthold et al. 1997, Nowald 2001). Their yearly return to breeding site, as well as their life-long pairing-up, make them particularly interesting for observations of climate effects and tracking of specific individuals. Due to their size, storks build large circular nests high up, in the open. Nests are thus quite sensible to meteorological variations, and the parental behavioral response to these variations is especially important for the chicks before the development of their thermoregulatory ability. On the other hand, passerines build well-insulated nests, with the use of feathers as material for tree swallows or mud for ovenbirds (Heenan 2013, Shibuya et al. 2015, Botero-Delgadillo et al. 2017). The orientation, with respect to the North, has been reported to be a criterium of selection for the birds, when choosing a nest box for instance or placing the entrance of an oven nest (Ardia et al. 2006, Schaaf and de la Peña 2018). For storks, the equivalent of the orientation of the nest is the orientation of the guarding parent. Yet, recent studies on stork nests have focused on the nest itself and show that such nests can accommodate a great diversity of invertebrates, plants and even passerines, suggesting that nest microclimates are also strong in stork nests (Zbyryt et al. 2017, Blonska et al. 2021, Dylewski et al. 2021). Few studies have looked into breeding correlation with weather conditions by storks (Tobolka et al. 2018, Kaminski et al. 2019). However parental behavior has yet to be considered in these studies, which is a point especially prone to modifications with climate changes (Durant et al. 2019, Mueller et al. 2019, Podkowa et al. 2019).

Our objective was first to understand the choice of placement of the stork parent on its nest, as preliminary observations suggested changes over the day. We predicted sun and wind to have significant influence on this choice. We thus correlated the orientation of the stork to the sun and wind direction, by combining image analysis of a webcam and meteorological data. Our study offers an overview of the parental care of a White stork expressed by its orientation and positioning on the nest during the nestling period of its chicks, together with some weather influence on this behavior.

## Material and methods

### Study species

The species taken for this study is white stork, *Ciconia ciconia.* The animals observed were a couple nesting on top of the city hall of Sarralbe, Moselle, France. This couple has been recorded to breed on this nest since 2016 with regular breeding success of about 3 fledglings per year. In 2020, they started incubation on March 15^th^ of 4 eggs. The third and fourth eggs were laid a couple of days later, as storks start their brooding on the laying of the second egg. Only three hatchlings were observed at the beginning of this study. They hatched after 32 days of incubation on April 16^th^ and the following days. Data recording and thorough observation started from May 2^nd^, time at which nestlings were 15, 17 and 18 days old. The age of the chicks is in the following considered to be the one of the youngest, so 15 days at the start of the observation, as well as the day reference is taken with the hatching of the latest chick.

### Data collection

The data was sampled through the observation webcam set on top of the city hall available to general audience (https://www.sarralbe.fr/Webcam.html). From this webcam, due to the impossibility of taking continuous recordings, captures were taken at various intervals from 5 minutes to 2 hours between 8am to 10pm, with a mean interval of 30 minutes. In total, 450 captures were done from May 2^nd^ to June 29^th^ with diminishing sampling rate due to both parents leaving the nest at the same time during longer time as nestlings grew up. Meteorological data (temperature, pressure and wind direction and intensity) were obtained *a posteriori* from the closest meteorological station, in Gros-Réderching, 20 km away from the nest. Sun position, represented by the angle *ψ* in Fig. 1 was computed from the geolocation of the nest and the hour of capture. The position of the stork on the nest was measured in polar coordinates (*l, θ*), taken at its feet, as shown in dark red in Fig. 1. The orientation of the stork, later referred to as *ϕ*, was calculated from the captures using MATLAB® image processing tools.

**Figure 1:**
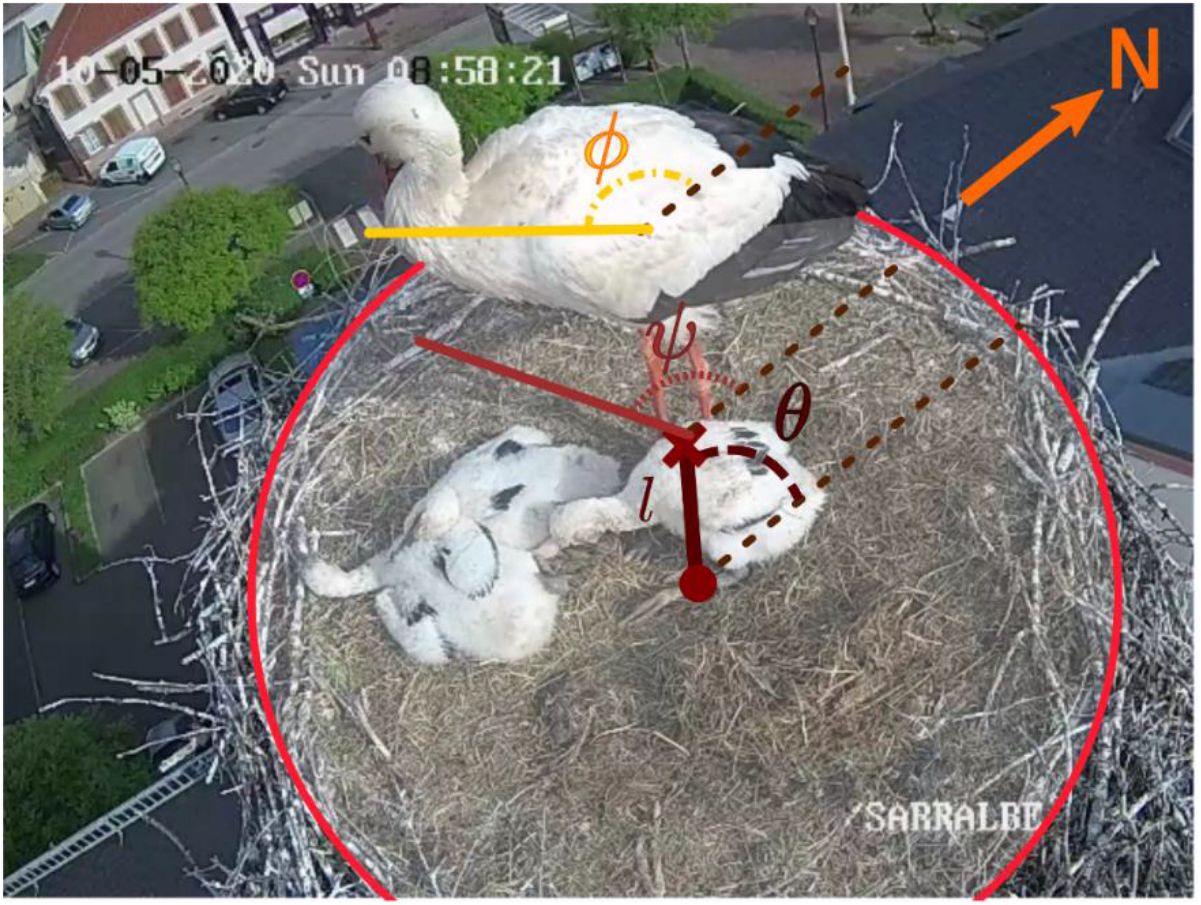
(colors online) sample image and measured angles (orientation in yellow and positioning, angle taken from the feet of the parent, in red). North is indicated in orange.

### Statistical analysis

To investigate the influence of weather on the choice in orientation of the guarding stork parent, I first looked into the overall occupation of the nest of the parent, through its positioning, i.e. distance to the center of the nest (*l*) and angular position (*θ*) with respect to the North. For this study, the age of the nestlings was a fixed factor while weather conditions were here used as random factors in first approximation. Then, weather impact was investigated on the orientation (*ϕ*) of the parent. As I could not ascertain normal distributions for the various parameters, I used Kruskal-Wallis tests instead of ANOVA tests for studying the influence of sun and time on the orientation. Temperature and wind were not found to be significant, except on one correlation as presented later. Wind strength and atmospheric pressure did not correlate with orientation and position. As such, they were excluded from the relevant parameters. No clear distinction was possible between the parents so sexual differences are not investigated in the following. Also, three observations of change in the guarding parent suggest that the choice of orientation is not individual-related as the new caring parent took each time the exact same position and orientation of its partner, that left the nest from a different position. Correlations were calculated using Spearman’s method. All statistical analysis was performed in the R environment version 4.0.3 (R core team 2020). Most interpretations come from graph-based considerations to complement statistical conclusions.

## Results

### Positioning on the nest

In Fig. 2.a, the occupation of the nest is represented over the whole duration of the study, one point corresponding to one sample. A first observation of the positioning of the stork parent on the nest is that there is no preferential location. There is a slight remoteness of the average position, the barycenter of all measured positions is indicated in light blue. Looking at the repartition in function of the age of the nestlings (coded in color), I observe that the younger the chicks, the closer the parent to the center of the nest. Angular position also seems to correlate with the age of the chicks. On the other hand, the hour of observation shows no correlation to the position, either with the distance to the nest nor with the angular position (see Table 1).

**Figure 2:**
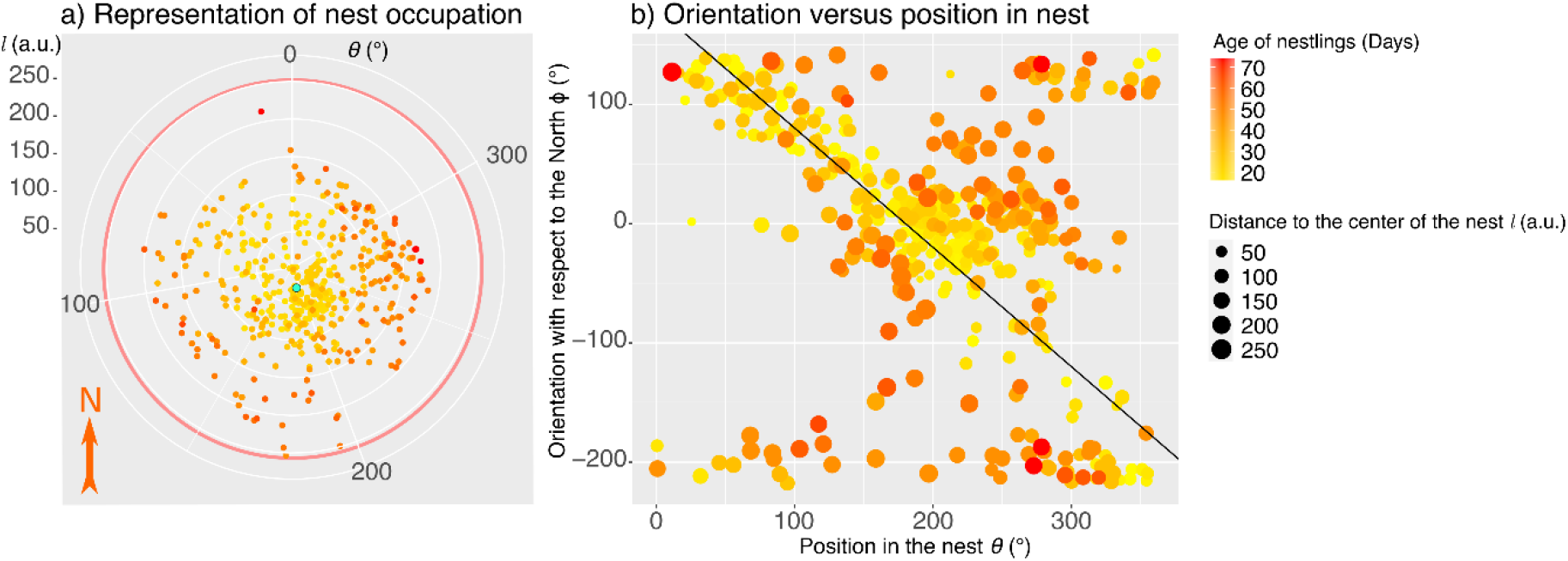
a) Nest representation with all measured positions of the stork parent over the observation. Edge of the nest is represented as in Fig. 1 by a red circle. b) Orientation of the parent with respect to its angular position in the nest. Black line represents the angles for which the sum of both is 180° (*ϕ* + *ψ* = 180°), corresponding to radial alignment of the stork. Color codes for the age of nestlings.

**Table 1:** Spearman’s correlations of age and observation time with position on the nest and orientation. Relevant correlations are stressed in bold.

| Correlation | | Distance to center $l$ | Angular position $\theta$ | Orientation $\phi$ |
| --- | --- | --- | --- | --- |
| **Day** (age of nestlings) | $r_s$ | **0.745** | **-0.17** | -0.06 |
| | $P$ | **<0.001** | **<0.001** | 0.25 |
| | $N$ | **450** | **450** | 450 |
| **Hour** (time of observation) | $r_s$ | -0.08 | 0.02 | **-0.17** |
| | $P$ | 0.07 | 0.68 | **<0.001** |
| | $N$ | 450 | 450 | **450** |

### Orientation in the nest

Once the position of the parent is analyzed, the orientation can be looked into as well and compared to the positioning. In Fig. 2.b, the orientation of the stork is presented against the angular position on the nest, with indications on the distance to the center on the size of the dots as well as the age of the chicks in color. Though the orientation looks dispersed at first glance, brighter data, thus in the early days of the chicks, are concentrated along a linear function in black. This function corresponds to the radial alignment of the stork in the nest. In the later days, multiple orientations are measured from the same position. When looking into the correlations to further investigate the relations observed in the figure, no correlation is observed between the orientation of the stork and the distance to the center. However, the angular position is strongly correlated especially up to around 35^th^ day of the nestlings. For observations before the 36^th^ day, the correlation coefficient is *r_s_* = 0.18 with *p* < 0.001, (*N* = 298) while for observations after the 34^th^ day, *r_s_* = −0.02 with *p* = 0.85, (*N* = 172).

### Effect of weather on orientation

The orientation and position being so intrinsically correlated, the question is now how weather conditions impact on the choice of either one. I separate these weather conditions into the sun presence (shadows visible in the image), its orientation (considered for every sample even when the sun is absent from the capture), the presence of rain (from meteorological data and observations of droplets in the image), the temperature and the wind direction *γ*. The sun orientation *ψ* is defined as follows: 0° meaning pointing toward the North. This can be also understood as the angle made between the shadow and the North, as shown in Fig. 1. As previously mentioned, the wind intensity and atmospheric pressure were not taken into account as relevant parameters.

Direct correlations between orientation of the stork and orientation of the sun shows a moderate effect size, *r_s_* = −0.175 with *p* < 0.001, *N* = 450 (see Table 1: hour is a proxy to the orientation of the sun). This correlation is even stronger when in presence of the sun (*r_s_* = −0.338 with *p* < 0.001, *N* = 246) and does not seem relevant when the sun is hidden behind clouds or set (*r_s_* = −0.077 with *p* = 0.277, *N* = 204). However, this simple correlation does not provide us with sufficient information on how the sun influences the choice of orientation. Therefore, I looked at the difference between the orientations of the stork and the sun with respect to the various parameters. In Fig. 3.a, this angular difference (in absolute value) |*ϕ* – *ψ*| is presented in regards to the absolute hourly time difference to the zenith (at 2pm in the region during that season). When the sun is present, |*ϕ* – *ψ*| is observed to be proportional to the relative time, especially below 4-hour difference. On the other hand, when the sun is absent, the angular difference is wide-spread and its average does not depend on the hour. This is supported by statistical analysis, through Spearman’s correlation, *r_s_* = 0.279 with *p* < 0.001, *N* = 450 in total, *r_s_* = 0.541 with *p* < 0.001, *N* = 246 for the sun present and *r_s_* = —0.017 with *p* = 0.809, *N* = 204 otherwise. The result of a Kruskal-Wallis test between both distributions further confirms a moderate effect with the total distribution, a large effect when the sun is present and a small effect when out. No effect of temperature is seen for the angle between Sun and Stork as correlation results for the three sets are *r_s_* ∈ [–0.04,0.03] with *p* > 0.5.

**Figure 3:**
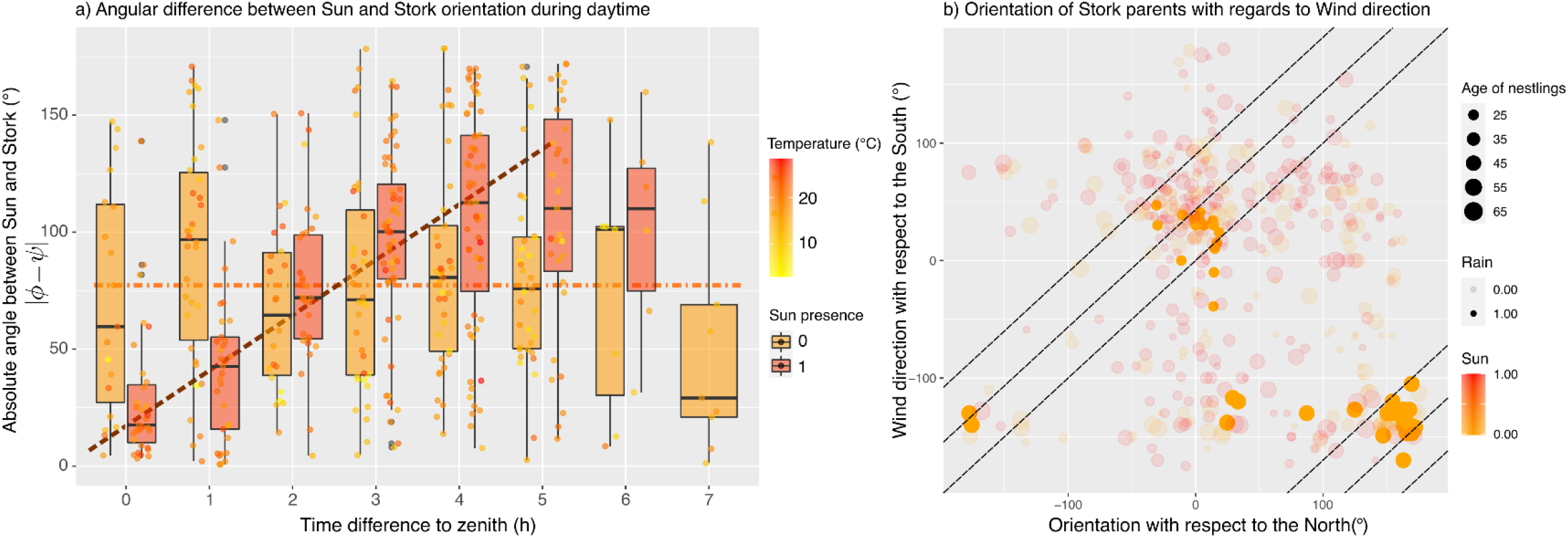
a) Difference between Sun and Stork orientations |*ϕ* – *ψ*|, as a function of the hourly difference to the zenith. Data are segregated by the sun presence. Dashed lines represent the overall tendency of evolution as visual guidelines. Color on the dots codes for temperature as indicator. b) Stork orientation (*ϕ*) vs wind direction (*γ*). Dashed lines represent the identity (*ϕ* = *γ*), and its translations *ϕ* = *γ* – 45° and *ϕ* = *γ* – 90°, for visual guidelines.

On the contrary, the wind direction (*γ*) does not show any correlation with the orientation when the sun is present, *r_s_* = 0.095 with *p* = 0.136, *N* = 246. Over all measurements, a moderate correlation is observed *r_s_* = 0.276 with *p* < 0.001, *N* = 450 . The highest correlation between wind direction and orientation is found when it rains, with *r_s_* = 0.715 with *p* < 0.001, *N* = 42. In Fig. 3.b, we see that the rainy data falls almost entirely around the −45° identity with a large confidence interval of 45°, *ϕ* = *γ* – 45°(±45°). On the other hand, measurements without rain are dispersed. The angle made by the stork with the wind |*ϕ* – *γ*| is correlated to the temperature, especially when it rains (*r_s_* = 0.702 with *p* < 0.001, *N* = 42, while almost uncorrelated when the sun is present (*r_s_* = 0.105 with p = 0.100, *N* = 246).

## Discussion

The results of this study demonstrate that neither the positioning nor the orientation of the stork are arbitrary but on the contrary carefully chosen depending on the weather conditions and the age of the nestlings. This conclusion of weather influence was hypothesized from the earliest observations of the nest ahead of quantitative measurements. It was further supported by the observations of change of brooding parent that took the exact same place and orientation as its pair as previously mentioned.

The initial result on the positioning being correlated to the age of the nestlings is understood by the fact that the stork parent leaves the center of the nest to its offspring for security. By gathering the chicks at the center, it is easier to guard them and the chicks have lower chances of falling from the nest. Yet as they grow, they occupy more space on the nest and the parent moves further away from the center. Their growth is also responsible for a change in behavior for the parents around 40 days old. The position on the nest no longer determines the orientation, or reverse, a single angular position may be used for multiple different orientations. In the early days, the position and the orientation coincide with the aspect of security guarding of the chicks. The stork parent is indeed in average constantly pointing toward the center of the nest. Once the chicks are old enough, not only do the parents leave them more often alone in the nest (Bochenski and Jerzak 2006, Duquet 2018) but I observe that when in the nest, they no longer orient toward the center. The change in behavior from the parents has been identified between the 36^th^ and 40^th^ day of the chicks, while in the literature, this change is observed around the 45^th^ day after hatching (Tortosa and Castro 2003, Tsachalidis et al. 2005, Duquet 2018). The 45^th^ day corresponds to the maximum of weight of the chicks but this limit can vary with weather conditions and food availability (Bochenski and Jerzak 2006). A way to identify nests with chicks past their maximum of weight could therefore be to look at the direction of the guarding parent and see whether it faces the center of the nest or not.

If the orientation is linked with the position solely due to the need of surveillance of the chicks, then, the question remains as to what decides on the position or on the orientation. I hypothesized that the orientation was the main parameter of choice for the stork parent, as I do not consider the stork to have preferable positioning with the support of the results (Fig. 2.a). With the need of thermoregulation for the nestlings, the sun and temperature were first thought to be determinant factors to the orientation. Yet the temperature did not influence the angle between the stork and the sun, which leads us to think that the orientation with the sun was not due solely to thermoregulation. The dependence however in the time to the zenith supports that the prime reason for the stork to orient is a balance between shadowing the brood and keeping its eyes protected from too much sun exposition. For shade provision, storks also use a *wing-drooping posture,* which enlarges their shadow while maintaining a safeguarding position (Bochenski and Jerzak 2006). With this, I conjecture that the orientation is chosen on sunny days from the sun direction and from its intensity or height in the sky, which proxy is the hour as it finds its maximum at the zenith, with the parent favoring wing-drooping posture and alignment to the sun when the latter would be too intense.

On the other hand, when the weather is inclement, with clouds or rain, the stork chooses its position from the wind. This may be understood from their dislike to having feathers brushed against. This is well-known in falconry where falconers always place themselves so that the bird faces the wind, as it turns toward it otherwise. Here, by the correlation between the temperature and the angle between the stork and the wind, this choice can also be interpreted by an issue in insulation and thermoregulation. Yet wind is reported to be of stronger impact when sleeking the feathers than fluffing it (Gebremedhin 1987, Bakken 1991), which would suggest that the stork would rather oppose the wind to minimize heat loss. These studies however did not account for rain and it is known by ducks that saturation in water in the feather can increase ten folds the heat loss (DeVries et al. 1995, Banta et al. 2004). Therefore, we can hypothesize that the stork avoids wind fluffing to prevent water penetration into the deeper coat, which would thus lead to further heat loss.

To conclude, the orientation of the parent stork on the nest is governed for the observed couple by the sun through its position in the sky and by the wind when the weather is inclement. Further observations coupled with meteorological data, from a station by the webcam for example, are required to draw statistically relevant conclusions for the species, also linking it to the breeding success, but this study offers a first insight on the question of orientation in the nest. Such studies, on weather influence on breeding success, have for instance recently been conducted for other species such as tits or finches (Gullett et al. 2015, Marques-Santos and Dingemanse 2020, Marques-Santos et al. 2021). Experiments in captivity would also be an asset to better understanding the role of wind for instance.

## Acknowledgments

I am thankful to the city of Sarralbe for the installation of the webcam on the nest, in particular Dominique Klein, who is the expert on stork and did most of the previous observations on this nest. I am grateful to my PhD supervisors, Nicolas Plihon and Mickaёl Bourgoin, for helping with the manuscript and allowing me to spend a bit of my research time on this project, despite being quite disconnected from my PhD work. Thanks also go to Maxen Cosset-Chéneau for fruitful discussions on preliminary observations.

## Bibliography

Ardia D. R., Pérez J. H., & Clotfelter E. D. 2006. Nest box orientation affects internal temperature and nest site selection by Tree Swallows. J. Field Ornithol. 77(3): 339–344.

Bakken G. S. 1991. Wind speed dependence of the overall thermal conductance of fur and feather insulation. J. Therm. Biol. 16(2):121–126.

Banta M. R., Lynott A. J., VanSant M. J., & Bakken G. S. 2004. Partitioning heat loss from mallard ducklings swimming on the air-water interface. J. Exp. Biol. 207(26):4551–4557.

Berthold P., van den Bossche W., Leshem Y., Kaatz C., Kaatz M., Nowak E., & Querner U. 1997. Satelliten-Telemetrie beim Weißstorch Ciconia ciconia: Wanderung eines Ost-Storchs westlich bis Nigeria. J. Ornithol. 138(3):331–334.

Blonska E., Lasota J. L., Jankowiak R., Michalcewicz J., Wojas T., Zbyryt A., & Ciach M. 2021. Biological and physicochemical properties of the nests of White Stork *Ciconia ciconia* reveal soil entirely formed, modified and maintained by birds. Sci. Total Environ. 763:143020.

Bochenski M., & Jerzak L. 2006. Behaviour of the White Stork Ciconia ciconia: a review. The White Stork in Poland: studies in biology, Ecol. Conserv. (January):295–324.

Bolton M., Butcher N., Sharpe F., Stevens D., & Fisher G. 2007. Remote monitoring of nests using digital camera technology. J. Field Ornithol. 78(2):213–220.

Botero-Delgadillo E., Orellana N., Serrano D., Poblete Y., & Vasquez R. A. 2017. Interpopulation variation in nest architecture in a secondary cavity-nesting bird suggests site-specific strategies to cope with heat loss and humidity. Auk 134(2):281–294.

Coates P. S., Connelly J. W., & Delehanty D. J. 2008. Predators of Greater Sage-Grouse nests identified by video monitoring. J. Field Ornithol. 79(4):421–428.

Duquet M. 2018. La cigogne blanche. Editions Delachaux et Niestlé, Lonay, Switzerland.

Durant S. E., Willson J. D., & Carroll R. B. 2019. Parental Effects and Climate Change: Will Avian Incubation Behavior Shield Embryos from Increasing Environmental Temperatures? Integr. Comp. Biol. 59(4):1068–1080.

Dylewski L., Dyderski M. K., Mackowiak, L. & Tobolka M., 2021. Nests of the white stork as suitable microsites for the colonisation and establishment of ruderal plants in the agricultural landscape. Plant Ecol. 222(3):337–348.

Gebremedhin K. G. 1987. Effect of animal orientation with respect to wind direction on convective heat loss. Agr. For. Meteorol. 40(2):199–206.

Gullett P. R., Hatchwell B. J., Robinson R. A., & Evans K. L. 2015. Breeding season weather determines long-tailed tit reproductive success through impacts on recruitment. J. Avian Biol. 46(5):441–451.

Halsey L. G., Portugal S. J., Smith J. A., Murn C. P., & Wilson R. P. 2009. Recording raptor behavior on the wing via accelerometry. J. Field Ornithol. 80(2): 171–177.

Heenan, C. B. 2013. An overview of the factors influencing the morphology and thermal properties of avian nests. Avian Biol. Res. 6(2):104–118.

Kaminski M., Banbura J., Janic B., Kaldma K., Konovalov A., Marszal L., Minias P., Vali U., & Zielinski P. 2019. Brood sex ratio and nestling physiological condition as indicators of the influence of weather conditions on breeding black storks *Ciconia nigra*. Ecol. Indic. 104(April):313–320.

Kumar N., Gupta U., Jhala Y. V., Qureshi Q., Gosler A. G., & Sergio F. 2020. GPS-telemetry unveils the regular high-elevation crossing of the Himalayas by a migratory raptor: implications for definition of a “Central Asian Flyway”. Sci. Rep. 10(1):1–9.

Laurent K., Fogg B., Ginsburg T., Halverson C., Lanzone M., Miller T., Winkler D. W., & Bewley G. P. 2021. Turbulence explains the accelerations of an eagle in natural flight, Proc. Natl. Acad. Sci. USA. 118(23):e2102588118.

Mandel J. T., Bildstein K. L., Bohrer G., & Winkler D. W. 2008. Movement ecology of migration in turkey vultures. P. Natl. Acad. Sci. USA. 105(49):19102–19107.

Marques-Santos F., & Dingemanse N. J. 2020. Weather effects on nestling survival of great tits vary according to the developmental stage. J. Avian Biol. 51:e02421.

Marques-Santos F., Wischhoff U., & Rodrigues M. 2021. Weather fluctuations are linked to nesting success and renesting decisions in saffron finches. J. Avian Biol. 52:e02323.

Mueller A. J., Miller K. D., & Bowers E. K. 2019. Nest microclimate during incubation affects posthatching development and parental care in wild birds. Sci. Rep. 9(1): 1–11.

Nowald G. 2001. Verhalten von Kranichfamilien (*Grus grus*) in Brutrevieren Nordostdeutsch-lands: Investition der Altvögel in ihre Nachkommen. J. Ornithol. 142(4):390–403.

Podkowa P., Malinowska K., & Surmacki A. 2019. Light affects parental provisioning behaviour in a cavity-nesting Passerine. J. Avian Biol. 50(11): 1–8.

Sabine J. B., Meyers J. M., & Schweitzer S. H. 2005. A simple, inexpensive video camera setup for the study of avian nest activity. J. Field Ornithol. 76(3):293–297.

Schaaf A. A., & de la Peña M. R. 2020. Bird nest orientation and local temperature: an analysis over three decades. Ecology 101(7):30–32.

Schulwitz S. E., Spurling D. P., Davis T. S., & McClure C. J. W. 2018. Webcams as an untapped opportunity to conduct citizen science: Six years of the American Kestrel Partnership’s Kestrel Cam. Glob. Ecol. Conserv. 15:e00434.

Shibuya F. L. S., Braga T. V., & Roper J. J. 2015. The Rufous Hornero (*Furnarius rufus*) nest as an incubation chamber. J. Therm. Biol. 47:7–12.

Stake M. M., Faaborg J., & Thompson F. R. 2004. Video identification of predators at Golden-cheeked Warbler nests. J. Field Ornithol. 75(4):337–344.

Thabethe V., & Downs C. T. 2018. Citizen science reveals widespread supplementary feeding of African woolly-necked storks in suburban areas of KwaZulu-Natal, South Africa. Urban Ecosystems 21(5):965–973.

Tobolka M., Dylewski L., Wozna J. T., & Zolnierowicz K. M. 2018. How weather conditions in non-breeding and breeding grounds affect the phenology and breeding abilities of white storks. Sci. Total Environ. 636:512–518.

Tortosa F. S., & Castro F. 2003. Development of thermoregulatory ability during ontogeny in the White Stork Ciconia ciconia. Ardeola 50(1):39–45.

Tryjanowski P., Kosicki J. Z., Kuzniak S. & Sparks T. H. 2009. Long-term changes and breeding success in relation to nesting structures used by the white stork, Ciconia ciconia. Ann. Zool. Fenn. 46(1):34–38.

Wood K. A., Ham P., Scales J., Wyeth E., & Rose P. E. 2020. Aggressive behavioural interactions between swans (*Cygnus spp.*) and other water birds during winter: A webcam-based study. Avian Res. 11(1):1–16.

Zbyryt A., Jakubas D., & Tobolka M. 2017. Factors determining presence of passerines breeding within White Stork *Ciconia ciconia* nests. Sci. Nat. 104(9-10):71.

